# Associations between sexual habits, menstrual hygiene practices, demographics and the vaginal microbiome as revealed by Bayesian network analysis

**DOI:** 10.1101/211631

**Authors:** Noelle Noyes, Kyu-Chul Cho, Jacques Ravel, Larry J. Forney, Zaid Abdo

## Abstract

The vaginal microbiome plays an influential role in several disease states in reproductive age women, including bacterial vaginosis (BV). While demographic characteristics are associated with differences in vaginal microbiome community structure, little is known about the influence of sexual and hygiene habits. Furthermore, associations between the vaginal microbiome and risk symptoms of bacterial vaginosis have not been fully elucidated. Using Bayesian network (BN) analysis of 16S rRNA gene sequence results, demographic and extensive questionnaire data, we describe both novel and previously documented associations between habits of women and their vaginal microbiome. The BN analysis approach shows promise in uncovering complex associations between disparate data types. Our findings based on this approach support published associations between specific microbiome members (e.g., *Eggerthella*, *Gardnerella*, *Dialister*, *Sneathia* and *Ruminococcaceae*), the Nugent score (a BV diagnostic) and vaginal pH (a risk symptom of BV). Additionally, we found that several microbiome members were directly connected to other risk symptoms of BV (such as vaginal discharge, odor, itch, irritation, and yeast infection) including *L. jensenii*, *Corynebacteria*, and *Proteobacteria*. No direct connections were found between the Nugent Score and risk symptoms of BV other than pH, indicating that the Nugent Score may not be the most useful criteria for assessment of clinical BV. We also found that demographics (i.e., age, ethnicity, previous pregnancy) were associated with the presence/absence of specific vaginal microbes. The resulting BN revealed several as-yet undocumented associations between birth control usage, menstrual hygiene practices and specific microbiome members. Many of these complex relationships were not identified using common analytical methods, i.e., ordination and PERMANOVA. While these associations require confirmatory follow-up study, our findings strongly suggest that future studies of the vaginal microbiome and vaginal pathologies should include detailed surveys of participants’ sanitary, sexual and birth control habits, as these can act as confounders in the relationship between the microbiome and disease. Although the BN approach is powerful in revealing complex associations within multidimensional datasets, the need in some cases to discretize the data for use in BN analysis can result in loss of information. Future research is required to alleviate such limitations in constructing BN networks. Large sample sizes are also required in order to allow for the incorporation of a large number of variables (nodes) into the BN, particularly when studying associations between metadata and the microbiome. We believe that this approach is of great value, complementing other methods, to further our understanding of complex associations characteristic of microbiome research.

## Introduction

The microbiome plays a critical role in human health, and the vaginal microbiome has been linked to urogential diseases of reproductive age women, including bacterial vaginosis (BV) (1–3). BV affects nearly one-third of women in the United States (4) and has been implicated in poorer pregnancy outcomes, acquisition of sexually-transmitted infections and other vaginal disorders (5,6). Specific changes in the vaginal microflora are associated with BV, including a depletion of *Lactobacillus* species and an increased abundance of strictly anaerobic bacteria (7). However, no single bacterial taxon has been shown to cause BV and the condition can be found in women with widely varying vaginal microbiomes (8–11). BV is characterized clinically by itching, pain, burning, odor and/or discharge, and is often diagnosed based on a combination of symptoms, vaginal pH and cytological findings (12). BV is also diagnosed using the Nugent score, which is the most commonly used diagnostic test for BV within the research community (13).

Just as the microbial underpinnings of BV are complex and varied (14), so too are the influences of a woman's sexual, sanitary and other practices. A wide range of factors have been shown to increase BV risk, including smoking, douching, menstruation, and new sexual partners (15–18). Different ethnic groups exhibit varying rates of BV, leading to conclusions that intrinsic host factors may contribute to the condition (4), although the effect of confounders on this association has been questioned (19). Further complicating this picture is evidence that women from different ethnic groups tend to harbor different vaginal microflora, independent of BV status (8,20).

BV is therefore a multifactorial disease mediated by a complex interplay of host, microbial and environmental factors. Fortunately, all three of these influences can be measured and evaluated using a combination of 16S rRNA gene sequencing of vaginal samples and detailed questionnaires of study participants. However, the analytical methods for identifying associations between multivariate, often categorical metadata with counts of microbial taxa are either ill-suited, contested or opaque (21,22). The most widely used analytical method for community-level data involves ordination of normalized microbiome counts (e.g., principal coordinates analysis [PCoA] and nonmetric multidimensional scaling [NMDS]) followed by statistical significance testing of explanatory/metadata variables (e.g., ANOSIM, PERMANOVA).

However, these methods reduce the dimensionality of the microbial community structure to two or three dimensions, obscuring intra-microbiome interactions. In addition, interactions between explanatory (metadata) variables are difficult or impossible to uncover because statistical testing usually occurs on a variable-by-variable basis (23). Therefore, these methods do not allow for discovery of more nuanced and complex dynamics between and within the microbiome and host and environmental factors. Identifying such dynamics can be crucial for understanding and thus combatting multifactorial conditions such as bacterial vaginosis – particularly when such conditions lack an accurate diagnostic description and/or test (24). Bayesian network (BN) analysis offers potential advantages in handling mixed datasets for complex conditions, and use of this approach may help to elucidate some of the nuanced, yet important, associations between host, environmental and microbial factors in diseases such as BV. For instance, BN's have been proposed as diagnostic tools for such multifactorial diseases as breast cancer and cardioembolic stroke (25–27). While the heavy computational burden of building BNs has impeded their widespread adoption, recent advances in algorithms have removed this barrier (28). In addition, software implementations now offer BN analysis within user-friendly packages (29,30). Finally, unlike other machine learning approaches, BN's possess an inherently intuitive interpretation and allow for a wide range of data input types, both continuous and categorical (31).

The primary goal of this study was to uncover associations between host behavioral characteristics, vaginal microbiome composition, risk symptoms and diagnostic criteria of BV utilizing BN's. Using this approach, we have demonstrated associations between women’s sexual and menstrual habits, demographics, vaginal microbiome composition, risk symptoms of BV and the Nugent Score (a BV diagnostic). Our findings support previously-documented associations between microbiome members (e.g., *Eggerthella*, *Gardnerella*, *Dialister*, *Sneathia* and *Ruminococcaceae*), the Nugent Score and vaginal pH (risk symptom) (8,13,32,33). However, we found no connections between the Nugent Score and other risk symptoms of BV such as vaginal discharge, itch, irritation, abdominal or pelvic pain, yeast infection and underwear staining within 60 days prior to sampling and any type of current vaginal odor. This suggests that the Nugent Score may not provide an accurate diagnostic of BV in some women. Additionally, we found that several microbiome members were directly connected to risk symptoms of BV, including *L. jensenii*, *Corynebacteria*, and *Proteobacteria*. Demographics (i.e., age, ethnicity, previous pregnancy) also influenced the presence/absence of specific vaginal microbes. The resulting network also revealed several as-yet undocumented associations between birth control usage, menstrual hygiene practices and microbiome members. These associations require confirmatory follow-up study, though strongly suggest that future studies of the vaginal microbiome and vaginal pathologies include detailed surveys of participants’ sanitary, sexual and birth control habits, as these can act as confounders in the relationship between the microbiome and disease.

Secondary objectives of this study included demonstrating the hypothesis-generating power of BN's through confirmation of previously described microbiome-BV associations, as well as the wide accessibility of BN's within a readily available, user-friendly environment. In addition, we aimed to illustrate the capacity of the BN method to directly associate component taxa of the microbiome with metadata. This capacity contrasts with common approaches used to analyze these data types (e.g., PCoA, NMDS and clustering), which rely on reducing the dimensionality of the total microbial community to two or three dimensions, followed by clustering or correlation of this reduced community with metadata (such as demographics). The BN approach also circumvents the limits on the number of interactions that can be tested between metadata and microbiome members that are imposed by tools such as PERMANOVA (34) or metagenomeSeq (35). This advantage of the BN approach is especially true when the number of variables observed in association with the microbiome is large. In such cases, network analysis identifies the hierarchy of relationships between the various metadata and microbiome taxa (i.e., the nodes of the network), and then represents these relationships using the arcs (or edges) of the network. When possible, we demonstrated these differences by providing direct comparisons of the BN approach with results obtained from Nonmetric Multidimensional Scaling Ordination (NMDS)-Analysis of Similarity (ANOSIM) and PERMANOVA. These comparisons should be interpreted with caution, as particularly the NMDS/ANOSIM approach is limited by the degrees of freedom available for significance testing and for assessing interactions. In addition -- and unlike the BN approach -- NMDS/ANOSIM and PERMANOVA analyses do not allow for the possibility of hierarchical relationships between variables observed within the data.

It is important to note that the BN approach presented in this work was aimed at inference, rather than prediction; and that we focused on evaluating associations within the data that could indicate possible associations in the target population. Use of the BN approach in this way does not attempt to classify the state of a new individual within the target population, i.e., prediction. Hence our network was fitted using all data and we did not attempt to perform cross validation to assess the ability of the network to predict or classify.

## Materials and Methods

### Study Population and Sampling

Questionnaire responses (see S1 File) and the relative abundance of 16S rRNA gene sequences (see S1 File) were obtained from a study of 396 healthy, sexually active, non-pregnant women of reproductive age (range 12-45 years); details of the study population have been described previously (8). Briefly, study participants submitted 2 self-collected vaginal swabs; one swab was used for 16S sequencing and the other was used to obtain a Nugent score, which is one of several diagnostic assays for BV and is the most commonly used within the research community (13). High Nugent scores are considered to be diagnostic of BV. Since all study participants were considered to be healthy, those with high Nugent scores were assumed to have asymptomatic BV. At the same time as vaginal swab collection, study participants were administered a detailed questionnaire on their sexual and sanitary habits and health histories (8, S2 File).

### Ethics Statement

The institutional review boards at Emory University School of Medicine, Grady Memorial Hospital and the University of Maryland School of Medicine approved the protocol. Guidelines of the universities were followed in the conduct of the clinical research. Participants provided written informed consent to participate in the study and written informed consent for use of the data for future studies. The study was registered at <u>clinicaltrials.gov</u> under ID NCT00576797.

### Data Preparation

Several filtering steps were taken with the goal of retaining only those variables (i.e., taxa, survey responses and demographic information) that would provide robust information during BN construction. In order to reach this goal, we removed taxa and survey response variables that were sparsely represented within the dataset, as described below.

Ribosomal 16S sequencing data were processed using the Ribosomal Database Project (RDP) Classifier (36) as described previously (8). In order to allow for robust BN construction, we removed samples that contained <1,000 16S counts. In addition, subjects who were not in overall general health or who had experienced toxic shock syndrome were removed from analysis. 16S counts were then normalized to the sample with the lowest number of counts, and taxa with counts <0.1% of total 16S counts were removed from further analysis to provide robust inputs into BN analysis. Finally, 16S counts were converted into presence/absence (i.e., 0/1) data (S3 File). While the implementation of BN analysis used in this study, i.e., bnlearn (29), is able to incorporate continuous variables with a Gaussian distribution, unfortunately the sparseness and skewness of the 16S rRNA count distributions in this dataset were such that we could not justify using abundances without dichotimization (see abundance distributions of used taxa in S8 File). Furthermore, categorizing these distributions to include more than presence/absence categories would have inflated the number of parameters that needed to be estimated, resulting in diminished stability of the BN network given the available sample size.

From the survey data, we removed questions that were related solely to study design factors (e.g., enrollment information, location of survey administration, survey administrator ID). Binary questions about specific vaginal odors (i.e., fishy, musty, foul and other) were collapsed into a single, binary "any odor" variable in order to decrease sparseness. In addition, we dropped questions about past birth control use in order to focus on current birth control habits, which we hypothesized would exhibit a stronger effect on the vaginal microbiome. Survey questions for which >5% of respondents did not provide an answer were excluded from BN analysis so that variables with a significant proportion of missing data did not exert undue influence over the graph. For the same reason, and after this variable-level exclusion, we removed any participants with missing data, keeping only those subjects with complete data in conjunction with the variables under study. After these filtering steps, we discretized any remaining continuous variables. Respondents' ages were converted from a continuous into a categorical variable with 3 levels: less than or equal to 30 (young adults), 31 – 40, and greater than 40 years of age (close to menopause). Nugent scores were categorized into low (0-3), medium (4-6) and high (7-10). Age at menstrual onset was discretized into <11, 11-15 and >15 years, and number of sexual partners in the last 60 days was categorized into 0, 1 and >1. Number of vaginal births and duration of menstruation were treated as categorical variables with 4 levels each (0 – 3 births and 1 – 4 days, respectively). Finally, we excluded survey questions with sparse outcomes, defined as having <5% of respondents within any one outcome level (i.e., for binary questions, at least 5% of respondents had to be in the "yes" and "no" outcome levels). This was also done such that stable BN's could be constructed. The resulting data are presented in S3 File and a key is available from S4 File.

### BN Construction

Filtered, dichotomized 16S rRNA gene counts as well as filtered and discretized demographic and survey responses were provided as input to BN construction, which was performed using bnlearn (29) implemented within the statistical software R (37). Age and ethnicity were specified as roots of the network, and Nugent score was specified as a leaf. Directionality between all other nodes was not specified in order to allow for the complex, likely multidirectional interplay between the microbiome, host factors, vaginal microenvironment and risk symptoms that might be associated with BV (12,14,38). The hill-climbing algorithm was used for BN construction with scoring based on the logarithm of the Bayesian Dirichlet equivalence (BDe) score; an optimal imaginary sample size was estimated using the alpha.star function in bnlearn (29,39). To obtain a consensus network, 1,000,000 BN's were constructed by first generating 1,000 random number seeds and, then, generating 1000 bootstraps of the input data. The aim of varying the seeds was to alleviate the possibility of being stuck in a local optimum in the space of all possible networks. The 1000 bootstraps per different starting point were used to assess stability of the network given each starting point. All 1,000,000 networks were then combined to construct a consensus BN by first using the mean function to pool the bootstrapped networks and then by using the averaged.network function to construct the consensus. Both of these functions are part of the bnlearn package. The empirical frequency for every arc within all BN's was determined, and arcs that met an empirical threshold frequency of 0.30 or more were used to produce the consensus BN. Network analysis and inference was performed using both this threshold and a 0.50 majority rule threshold. The resulting DAG was used in maximum likelihood estimation parameter fitting through the bn.fit function, as well as network analysis (29).

### BN Analysis

Prior to BN analysis, nodes were categorized as either “demographic”, “microbiome”, “sexual/sanitary habit”, “BV risk symptom”, or “BV diagnostic criteria” (S4 File). Variables that were categorized as BV risk symptoms included “staining of underwear”, “vaginal discharge”, “vaginal odor”, “vaginal itch”, “vaginal irritation”, “abdominal or pelvic pain” and “yeast infection” within the past 60 days prior to sampling, and pH and any type of current vaginal odor (any odor). These variables where chosen based on previously observed associations with increased risk of BV (6,12,40). Vaginal pH, vaginal discharge and staining, and odor could be thought of as proxies for the Amsel symptoms used in BV diagnosis in clinical settings (32). Given that all included subjects were considered healthy at the time of this study, our intention was to highlight possible increased risk of BV due to presence of these risk symptoms. The Nugent Score was the only variable designated as a “BV diagnostic criteria”. After *a priori* categorization, the nodes and edges obtained from bnlearn were used to conduct network analysis in order to understand the structure of the overall network, as well as the connections within it. Average Markov blanket size and overall graph characteristics were calculated and reported. Network density was defined as the proportion of all possible edges that occurred in the final network. Node degree was defined as the number of incoming and outgoing edges, node closeness centrality was defined as the average shortest path from a node to all other nodes, and node betweenness centrality was defined as the frequency with which a node was included in the shortest path between nodes in the network. Node closeness and betweenness centralities were normalized for comparison purposes. Groupings of nodes (i.e., groupings not related to the *a priori* node category, but rather agnostic groupings) were detected using a modularity optimizing algorithm (41) as implemented in Gephi (42), using a resolution of 1.0 and randomization without edge weights. Networks were visualized using Gephi (42), using the Force Atlas layout unless otherwise specified.

### Ordination and Clustering

Filtered, normalized, non-dichotomized, Hellinger-transformed (43) 16S rRNA gene counts were used to construct a Euclidean distance matrix, which was then ordinated using NMDS with multiple random starts and two dimensions (44). Use of two dimensions resulted in a stress value of 0.19, indicating less-than-ideal fit; however, increasing the number of dimensions resulted in failure of convergence, and therefore a better fit could not be achieved. Associations between the ordinated microbiome and metadata variables (i.e., host demographics and survey data) were tested using analysis of similarities (ANOSIM) (45). A *P*-value of less than 0.05 was considered statistically significant.

### PERMANOVA

Permutation based multivariate analysis of variance (PERMANOVA) was also used to assess the influence of metadata on the vaginal microbiome. The process of assessing significance of association followed a forward elimination, stepwise model selection approach based on Akaike’s Information Criteria (AIC) and using the step function as implemented in R. PERMANOVA was performed onHellinger-transformed microbiome counts. All survey and demographic response variables that passed the stepwise selection criteria (see above) and minimized the AIC score were included in the final PERMANOVA model (implemented using rda function in vegan). A permutation test using 1000 permutations was used to compute *P*-values to assess significance of associations between the metadata and the observed microbiome.

## Results and Discussion

### Data filtering and descriptive statistics

Of the 396 participants included in the original study, 2 were removed *a priori* due to a lack of any metadata from the questionnaire. Among the remaining 394 participants, 247 genus- and species-level taxa were identified through 16S sequencing of vaginal swabs. Total 16S counts per sample ranged from 693 to 7,392 with a median of 2,149. Five samples with <1,000 total 16S counts were removed from the analysis, for a remaining 389 samples; all 247 taxa were represented within these 389 samples. 16S counts were normalized to a count of 1,006, which was the total number of 16S reads in the sample with the lowest total count. The distribution of 16S counts was strongly right-skewed, and therefore 220 taxa were present at <0.1% abundance (i.e., fewer than an average of 2 reads per sample) and were removed from further analysis. From the 389 study participants with 16S data that passed filtering, a further 4 participants were removed because they answered "no" or did not answer the question of whether they were in overall good health; and a further 9 were removed because they either answered "yes" to previous toxic shock syndrome or did not answer this question. A further 91 respondents were excluded from analysis due to missing data for survey questions. The decision to remove subjects with missing data was not made lightly and was based on two reasons: 1) data imputation, although useful, could be faulty or could reduce the variability of the estimates of the network parameters (46) resulting in overconfidence in the network and 2) adding the missing data as a category by itself would substantially hamper interpretability of results. Accordingly, the decision was made to err on the conservative side (i.e., increased variability), rather than on the side of overconfidence or weak interpretability. This left a total of 285 healthy study participants for inclusion in BN analysis, encompassing a total of 27 taxa with a median prevalence of 89/285 samples (31%, range 24 - 228 out of 285 samples, S3 File). Among the 285 participants included in the network, 58 self-reported to be Hispanic (20.4%), 71 Asian (24.9%), 78 White (27.4%) and 78 African-American (27.4%). Nearly half of respondents were age 30 or under (49.5%, 141/285), and only 25 reported being over age 40 (8.8%).

After removing study-specific questions, redundant questions, past birth control use questions, questions about frequency of sexual habits and menstrual hygiene, as well as questions related to study exclusion criteria, a total of 263 survey questions were included in the initial survey response data. After removing questions with >5% missing responses, we were left with 56 study questions. A further 27 questions were removed due to sparseness, resulting in a total of 29 study questions available for input into the BN (S3 File). A majority of the 285 survey respondents included in the Bayesian network reported having been pregnant at least once (170/285, 59.6%). Sixty-three reported having had no sex partners in the 60 days preceding sampling (22.1%), a further 199 were monogamous in this same time period (69.8%), and 23 respondents had multiple sex partners (8.1%). The majority of women reported an absence of vaginal itching, irritation, discharge, odor or uro-abdominal pain in the 60 days preceding the survey (177/285, 62.1%). Vaginal yeast infections in the 60 days preceding sampling were reported by 19 of the 285 respondents. In addition to the 29 survey questions, we also had data on the vaginal pH and Nugent score of the 285 women in the study. The majority of participants had a low Nugent score (188/285, 66.0%), while a smaller proportion were categorized as intermediate and high (34/285 and 63/285, 11.9% and 22.1%, respectively). Around half of the 285 women had a vaginal pH of less than or equal to 4.5 (152/285, 53.3%), while about half had a higher vaginal pH (133/285, 46.7%).

### Network construction

A total of 60 variables were included in BN construction: the 29 survey response variables and 27 bacterial taxa that passed filtering criteria, as well as age, ethnicity, vaginal pH and Nugent score. The consensus BN had 152 directed edges with an average Markov blanket size of 9.4.

### BN Characteristics

The final consensus graph contained 60 nodes within a single sparse network with a density of 0.043. The majority of nodes contained between 1 and 6 connections, while several were more highly connected (Fig. 1). *Eggerthella* represented the most highly-connected node with 11 edges, followed by *Sneathia*, previous pregnancy status and current use of any type of birth control with 10 edges each.

**Figure 1.**
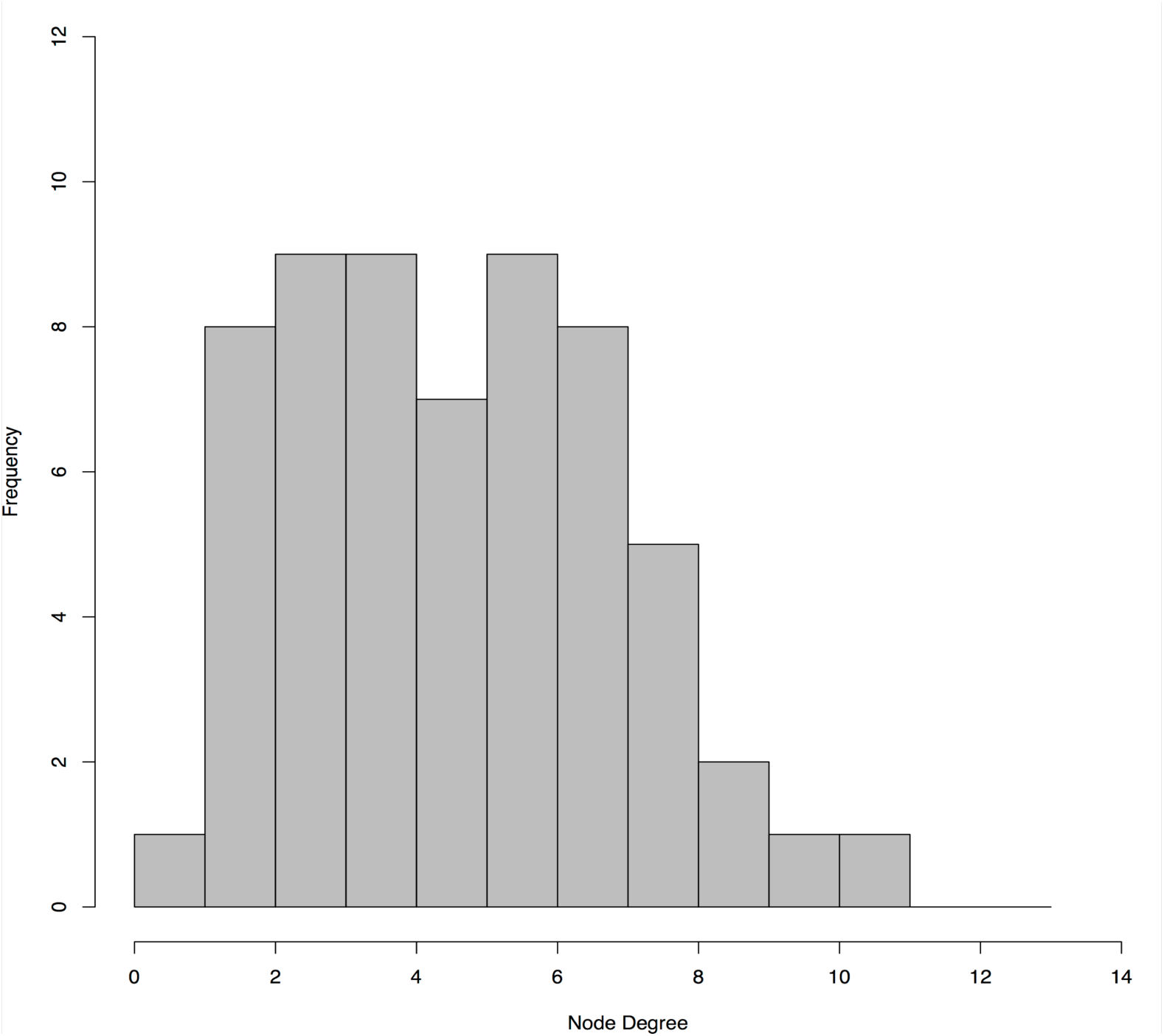
Histogram of node degree for the final BN. The majority of nodes had fewer than 8 connections (i.e., degrees), while a few were more highly connected.

Betweenness centrality measures indicated that *Parvimonas*, report of vaginal odor in the 60 days prior to the survey, as well as presence/absence of *L. vaginalis* within the vaginal microbiome, were situated along highly-connected paths in the network [S4 File, see column “Betweenness Centrality”]. Node closeness centrality metrics suggested that presence/absence of *Lactobacillus iners, Mycoplasmataceae, Lachnospiraceae* and *Streptococcus* were all nodes at the geographic center of the network, along with vaginal irritation in the 60 days preceding the survey [S4 File, see column “Closeness Centrality”]. Interestingly, *Sneathia*, *Eggerthella*, *Parvimonas* and *Streptococcus* have been described as key members of a vaginal microbiome characterized by a diversity of non-*Lactobacillus* bacteria found more commonly in black and Hispanic women (8,47). The fact that these bacteria are centrally situated in the BN developed in this study (as defined by closeness, betweenness and degree metrics) lends support to the idea that these bacteria could play an important role in differentiating the vaginal microbiome in different women.

### Ordination, Clustering and PERMANOVA

Univariable clustering analysis with ANOSIM significance testing revealed that six of the 33 metadata variables (including host demographics and survey responses) were statistically significantly associated with the microbiome ordination results. This list comprised ethnicity, vaginal pH, Nugent score, tampon use, as well as previous pregnancy and vaginal birth statuses [S5 File]. S6 File provides the associated NMDS plots for these six variables. These analyses suggested that demographic and behavioral traits of participants were associated with differences in the microbiome; however, given the nature of these analyses, it was unclear specifically which microbiome members were being affected, and whether such associations were direct or indirect. PERMANOVA analysis, alternatively, revealed a best-fit model that also included vaginal pH, Nugent score and previous pregnancy, as well as participant age, use of pads during menstruation, staining of underwear in the previous 60 days, and use of male condoms as current birth control method, which together explained 25% of the variability in the data (Table 1). This type of analysis provided more information highlighting associations between high Nugent Score, *Sneathia* and *Megasphaera* and possibly *Atopobium*, *Dialister*, *Eggerthella*, *Ruminococcaceae*, and *Lachnospiraceae* (S7 Figure). It also showed putative association between intermediate Nugent Score and *L. gasseri* (S7 Figure). Other relationships were tenuous and not clearly defined. Furthermore, the above analyses precluded the testing of more complicated interactions between multiple variables of diverse types. BN analysis was thus used to identify such relationships within all of the data, including between individual microbiome members; we point to the results of NMDS/ANOSIM and PERMANOVA where relevant.

**Table 1.** PERMANOVA permutation test results after stepwise model selection using the AIC. PERMANOVA was used to assess the significance of associations between metadata and the vaginal microbiome composition. P-values are based on 1000 permutations.

| **Variable** | **Degrees of freedom** | **p-values** |
| --- | --- | --- |
| **Nugent score** | **2** | **0.001** |
| **Previous pregnancy** | **1** | **0.001** |
| **Vaginal pH** | **1** | **0.001** |
| **Participant age** | **2** | **0.012** |
| **Use of menstrual pads during menstruation** | **1** | **0.023** |
| **Staining of underwear in previous 60 days** | **1** | **0.027** |
| **Use of male condoms as current birth control method** | **1** | **0.066** |

### Associations between population demographics, the microbiome and survey responses

There were 58 edges that bridged nodes of different types, i.e., demographics, sexual and menstrual hygiene habits, risk symptoms of BV, diagnostic criteria and microbiome (see S4 File for *a priori* classification of nodes); compared to 94 edges that connected nodes of like types. Of these 58 bridging edges, 16 were identified in 50% of the bootstrapped BN networks (Fig. 2). Many of these connections confirmed previously documented relationships between demographic factors and sexual and menstrual habits. For instance, ethnicity was directly related to Pap testing status, with Asian women exhibiting a much higher likelihood of never having had a pap smear compared to Hispanic, White and Black respondents (20.8% versus 4.6%, 2.9%, and 1.2%, respectively), a disparity that has been described previously (48,49). At a threshold of at least 50% bootstrap support, ethnicity also influenced tampon and pad use during menstruation (50), while at 30% support ethnicity was associated with age at menstrual onset (51,52) and the likelihood of a woman ever having self-treated a vaginal infection.

**Figure 2.**
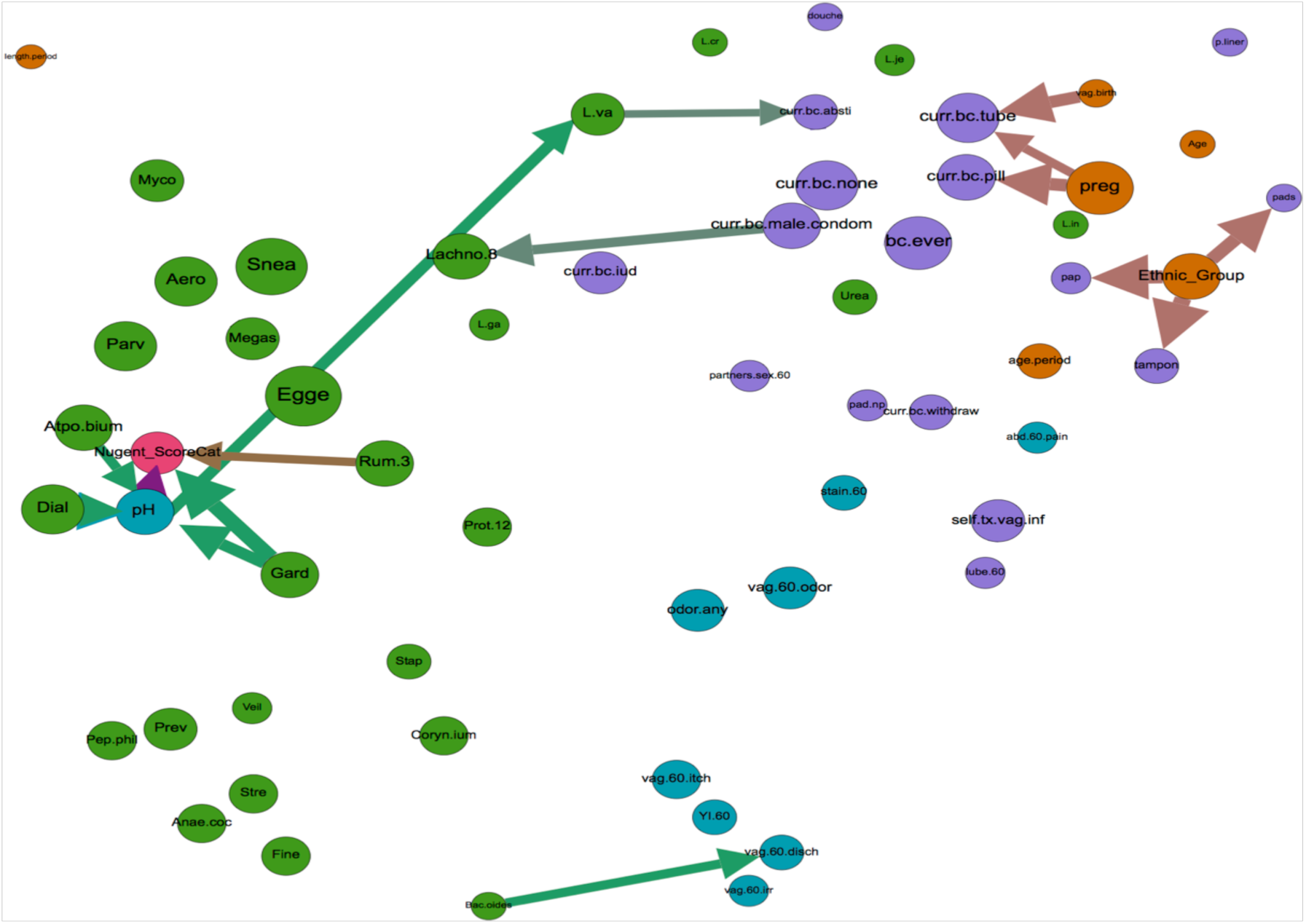
Inter-category bridging edges with >50% bootstrap. Network displaying all nodes, but depicting only inter-category bridging edges with >50% bootstrap support (arrows). Node size is proportional to closeness centrality, arrow thickness is proportional to bootstrap support, and node color signifies category type (orange = demographic, green = microbiome, turquoise = BV risk symptom, purple = sexual/menstrual habits, and pink = diagnostic criteria of BV). Node label abbreviations and categorization by type can be found in S4 File.

At 50% bootstrap support, there were no edges between demographic variables (i.e., age, ethnicity, age at menstrual onset, previous pregnancy and vaginal birth) and microbiome nodes (Fig. 2), suggesting that these factors may not exert robust influence over the presence of common microbiome members. However, decreasing bootstrap support to 30% revealed four such edges, namely associations between participant age and *Lactobacillus iners*, ethnicity and *Eggerthella*, age at menstrual onset and *Ureaplasma*, and *Mycoplasmataceae* and period length (Fig. 3). PERMANOVA testing supported the associations between the vaginal microbiome and age, while NMDS/ANOSIM analysis supported the association between the microbiome and ethnicity and marginally (ANOSIM *P* = 0.06) between the microbiome and age at menarch (Table 1, S5-S7 Files). Additionally, PERMANOVA and NMDS/ANOSIM analyses revealed associations between the vaginal microbiome and previous pregnancy and/or previous live birth (Table 1, S5-S7 Files). Given the constraints of PERMANOVA and NMDS/ANOSIM analysis, we were unable to identify which specific microbiome members may be involved in these associations.

**Figure 3.**
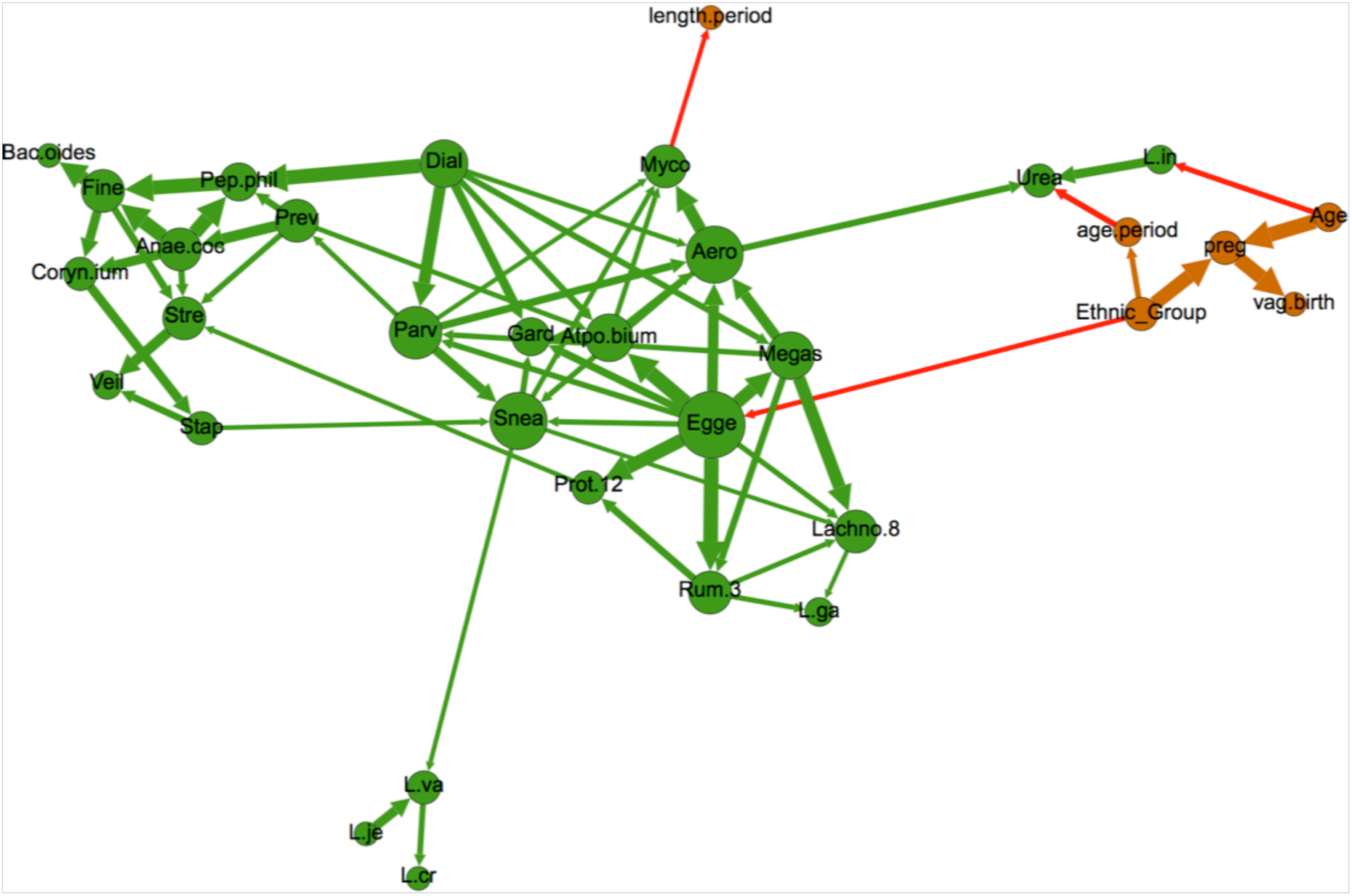
Network depicting nodes related to demographics (orange) and microbial taxa (green), as well as all edges with at least 30% bootstrap support. Node size is proportional to node degree (i.e., number of incoming and outgoing edges). Arrow thickness is proportional to bootstrap support. Red arrows bridge demographic and microbiome nodes, while green and orange arrows connect nodes of the same type (i.e., microbiome-to-microbiome or demographic-to-demographic). Node label abbreviations can be found in S4 File.

However, BN analysis revealed that the probability of harboring *L. iners* given age >40 years was 93%, compared to 72% and 85% for women ages 30-40 and <30 years, respectively. To date, longitudinal studies of the vaginal microbiome have been restricted to relatively short time periods, with subsequently little knowledge about how the microbiome may change with age. However, studies suggest that variations in estrogen levels influence presence/abundance of *Lactobacilli* (53,54), and age has a significant influence on estrogen levels (55), which together present a possible mechanism for the association we found between age and *L. iners*. Existence of an edge between these two nodes suggests significance of association (affirmed by a Chi-square test-of-independence, p-value 0.0067). BN analysis also revealed a complex relationship between age and other demographic variables. Age was associated with previous pregnancy status (which can also affect estrogen levels), as was ethnicity; indeed, previous pregnancy was a common effect of age and ethnicity within this dataset (i.e., the three nodes formed a v-structure within the graph). Only 24% of white respondents reported having ever been pregnant (19/78), compared to 58%, 77% and 86% of Asians, African-Americans and Hispanics, respectively. The role of age in likelihood of previous pregnancy was very strong among Asian and white respondents, and less so for African-Americans and Hispanics. Ethnicity also influenced the likelihood of identifying *Eggerthella* within the microbiome, as African-Americans were more likely to harbor this bacterium (45.4% versus 18.8%, 16.5% and 3.8% for Hispanics, Caucasians and Asians, respectively). Interestingly, *Eggerthella* was one of 28 taxa that were previously found to exhibit a significant interaction with race and BV status (47), and was also a key member of a certain vaginal microbiome composition that was more commonly found in black and Hispanic women than white and Asian women (8). Finally, previous pregnancy decreased the likelihood of a woman harboring *L. crispatus* and age at menarch was associated with *Ureasplasma* presence. *Ureaplasma* presence was also influenced by pap smear status. *Ureaplasma* in the upper genital tract has been associated with poor pregnancy outcomes, and *Ureaplasma* colonization in the lower genital tract has been found to be associated with a variety of socioeconomic and demographic factors including educational, income, ethnicity and marital status (56–58). Interestingly, age at menarch has also been associated with socioeconomic-related factors such as body mass index (BMI) and ethnicity (59,60), as has pap smear status (48,49). Results such as these demonstrate the complexity of interactions between microbiome composition and demographic and behavioral factors, and strongly suggest that any attempts to associate microbial composition with clinical BV should include co-analysis of potential confounders. The results presented here indicate that age, ethnicity and previous pregnancy status could potentially be used as portmanteau variables for such confounders.

Upon modularity analysis, nodes within the network tended to segregate into groupings of nodes of like type (called “communities” in network analysis), with some exceptions (Fig. 4). Nearly all of the microbiome nodes fell into two largely bacteria-specific groupings, one of which contained the nodes for Nugent score and vaginal pH. All of the *Lactobacilli* except *L. gasseri* segregated along with nodes related to demographics (including study participant age, ethnicity, pap smear status, age at menstrual onset and previous pregnancy), menstrual hygiene habits and douching. All of the nodes related to risk symptoms associated with increased risk of BV, except for pH, fell into one grouping that also included use of the withdrawal method for birth control, and all of the other birth control variables were clustered into a fifth grouping that also included the number of sexual partners in the previous 60 days (Fig. 4).

**Figure 4:**
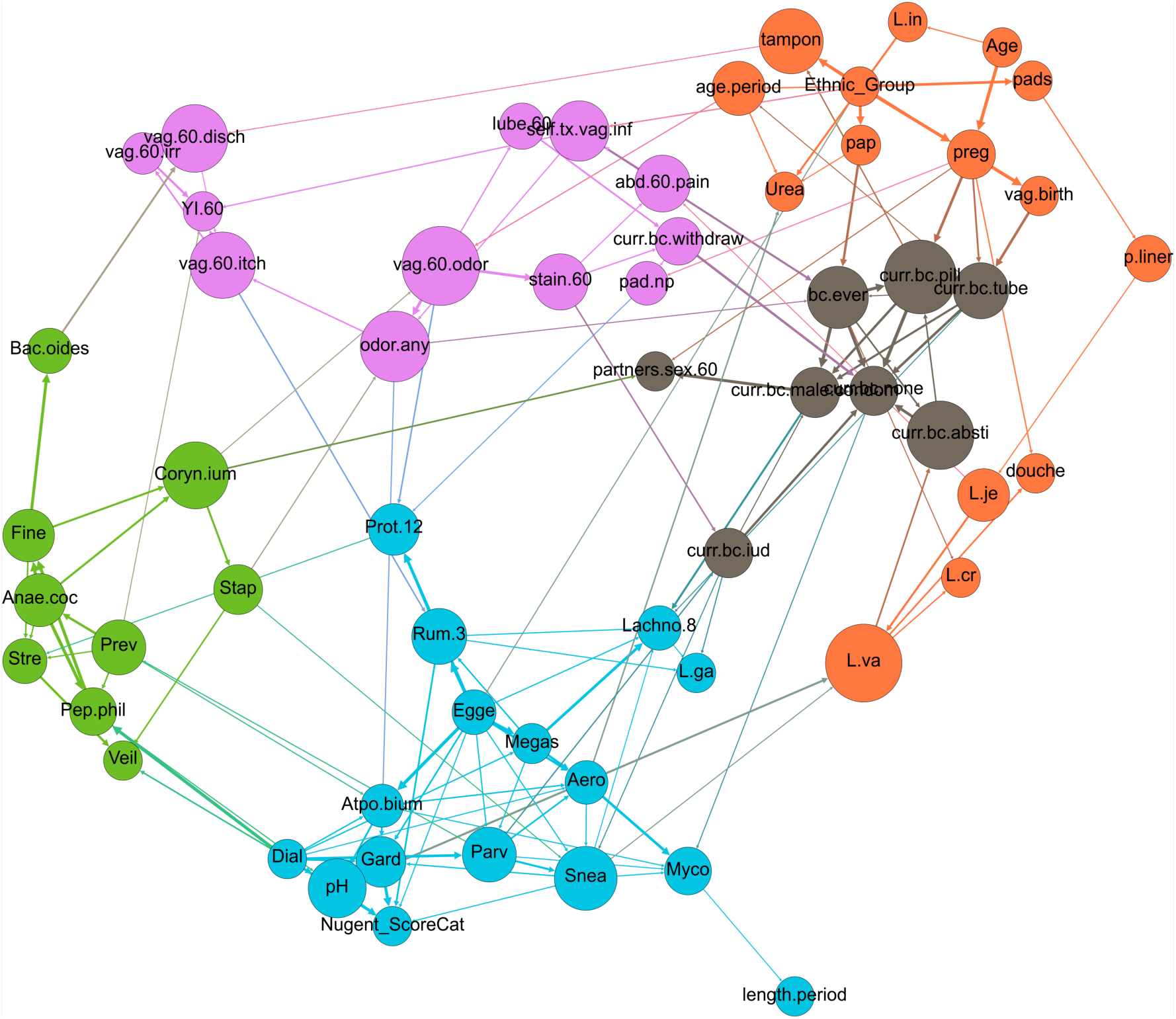
Network depicting modularity-based communities (node colors) as determined by a modularity optimization algorithm (41). Size of node is proportional to node’s betweenness centrality. Node label abbreviations can be found in S4 File.

These modularity-based grouping results highlight the dissociation of the Nugent score (a BV diagnostic) and vaginal pH from all other risk symptoms of BV, a finding which supports recent studies suggesting that these may not be the most useful criteria for assessment of clinical BV disease (8,33). None of the other risk symptom-based nodes fell into the same modularity grouping as the Nugent score and vaginal pH, and indeed the BN contained no directed paths of any length between vaginal pH or Nugent score and risk symptoms of vaginal irritation, itch, odor, pain or staining. Furthermore, contingency table propagation showed that women reporting vaginal odor, itch and irritation had an 11.6% chance of a high Nugent score, compared to 11.0% for women who did not report vaginal odor, itch and irritation, again suggesting that the Nugent score might not be reliably associated with these risk symptoms. In addition, while bacteria traditionally thought to be determinants of the Nugent score did belong to the same modularity grouping as the node for Nugent score (e.g., *Dialister* and *Gardnerella*), these bacteria were not associated with BV risk symptoms other than pH (Fig. 4).

### Associations between pH, Nugent Score and the microbiome

Several microbial taxa nodes were directly associated with vaginal pH and the Nugent Score. Vaginal pH is one of the four Amsel criteria used to diagnose BV (32). A vaginal pH of less than or equal to 4.5 is considered normal, while >4.5 is considered to contribute to BV. Although all women in this dataset were healthy and did not have clinical BV as determined by the Amsel criteria, nearly half had an “abnormal” vaginal pH >4.5 (133/285, 46.7%), a finding which calls into question the use of vaginal pH as an indicator of clinical BV. As with previous studies (61), NMDS/ANOSIM and PERMANOVA analyses both showed association between the vaginal microbiome and vaginal pH (Table 1, S5-S7 Files). Using BN analysis, we were able to reveal that vaginal pH was influenced by 4 bacteria: *Atopobium*, *Dialister, Gardnerella* and *Peptoniphilus* (Fig. 5). These bacteria are all obligate or facultative anaerobes and the presence of each of these bacteria in the vaginal microbiome increased the likelihood of a vaginal pH > 4.5. These four bacteria have been described as key members of a vaginal microbiome community type characterized by a diversity of non-*Lactobacilli* bacteria that is found more commonly in women with a high vaginal pH. It has been hypothesized that the higher vaginal pH in such women is due to a comparatively low number of lactic acid producing bacteria (8), and the results of this study support this hypothesis. As with previous research, however, the presence of these bacteria and the subsequent increase in vaginal pH were reported in healthy women without clinical BV.

**Figure 5.**
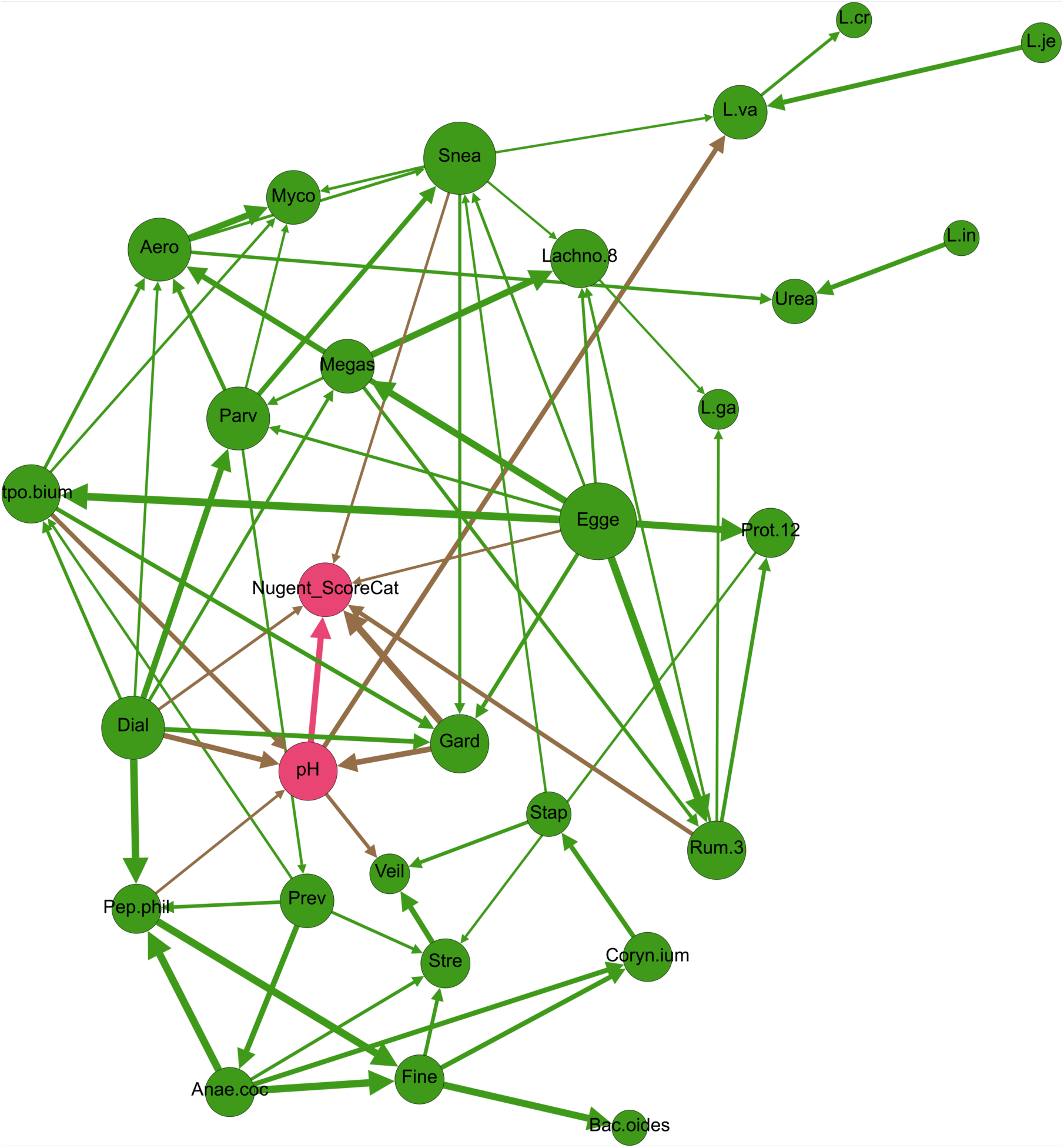
Network depicting vaginal pH and Nugent Score (pink), and bacteria (green). Size of circle is proportional to degree (i.e., number of incoming and outgoing edges), and edge thickness is proportional to bootstrap support. Node label abbreviations can be found in S4 File.

The Nugent score is a diagnostic test used primarily in research studies of BV (13). It is rarely used in practice because it involves time-consuming microscopic examination of bacterial morphology from vaginal swabs. The Nugent score is based on the relative presence of Gram-positive versus Gram-negative straight and curved rods, and therefore it is unsurprising that NMDS/ANOSIM and PERMANOVA testing both uncovered associations between the vaginal microbiome and the Nugent score (Table 1, S5-S7 Files). Using BN analysis, we were able to assess this association more closely in order to uncover which specific microbiome members were driving differences in the Nugent score. We found that the Nugent score for the 285 women in this dataset was influenced by bacteria from two Gram-positive bacteria (*Eggerthella* and *Ruminococcaceae*), two Gram-negative bacteria (*Dialister* and *Sneathia*) and one Gram-variable bacterium (*Gardnerella*) group. Vaginal pH was also connected to Nugent score; women with a pH of >4.5 were much more likely to have a high Nugent score, a finding consistent with previous literature (8,62). Amongst the 64 combinations of parent variable states that directly influenced the Nugent Score, there were 6 combinations that conferred 100% conditional probability of a high Nugent score; all 6 combinations included *Dialister*, *Ruminococcaceae* and *Sneathia*, which have been found in higher abundance in women with a high Nugent score (8, Fig. 6). Overall, these results confirm previous findings concerning the robust association between certain vaginal microbiome members and vaginal pH and Nugent score. However, the women in this study were BV-negative, and therefore the presence of high vaginal pH, a high Nugent score and/or microbes associated with these indicators does not always correlate with clinical BV (8). It is worth noting that the *Lactobacillus* species and the Nugent Score were only conditionally independent given the presence of vaginal pH and *Sneathia—*which were directly associated with *L. vaginalis* and indirectly with both *L. jensenii*and *L. crispatus*—and *Ruminococcacea*, which shared a direct edge with *L. gasseri*. This indicates a weaker association between the Nugent Score and the *Lactobacilli*, which was masked by the presence of stronger associations, e.g., with vaginal pH. Removing some of these directly-linked nodes from the network resulted in a direct association of the Nugent score with some *Lactobacillus* nodes (data not shown).

**Figure 6.**
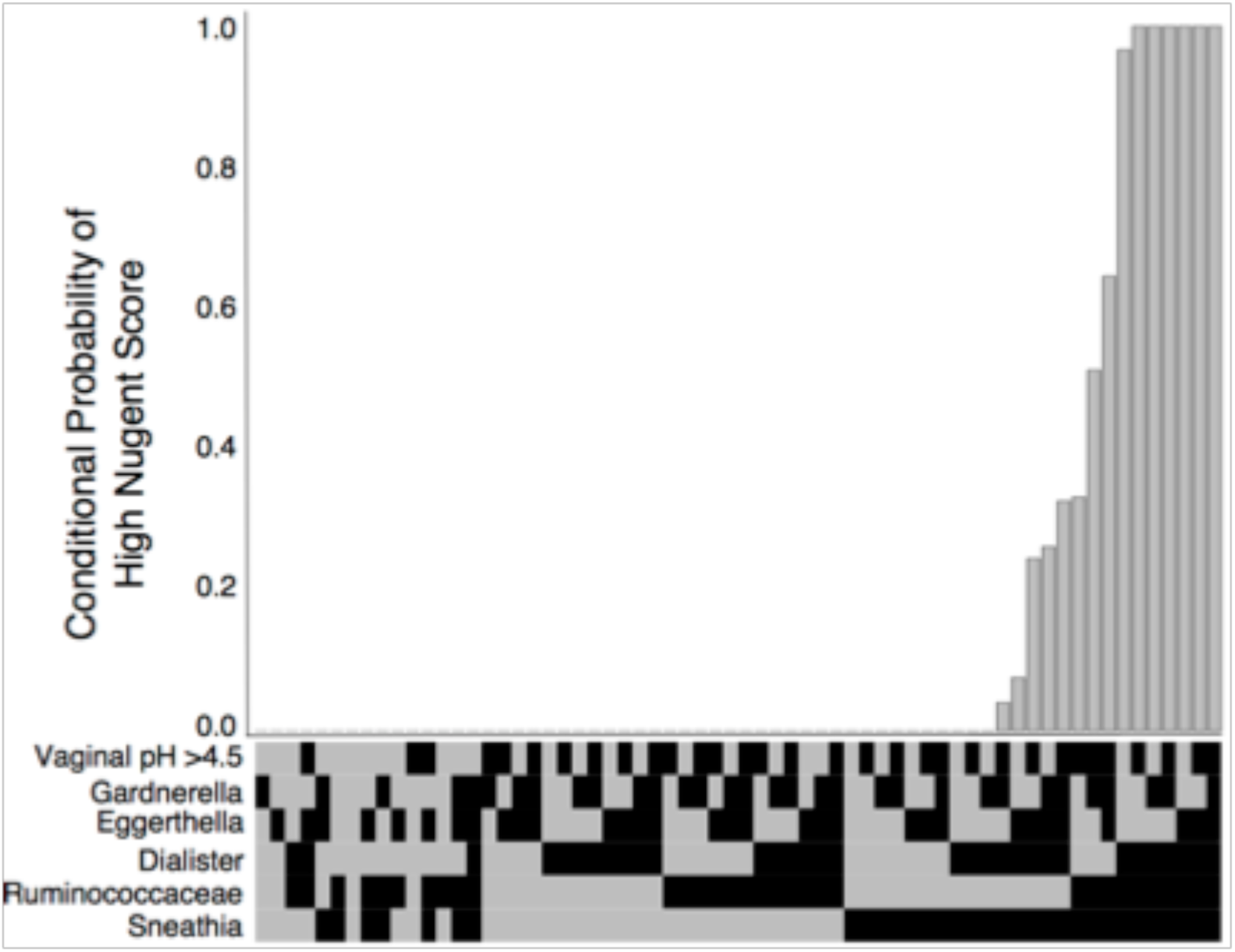
Conditional probability of a high Nugent Score (y-axis) for all possible combinations of direct parent variables (x-axis) (binary heatmap, black = present, gray = absent).

### Associations between the microbiome and risk symptoms other than vaginal pH

While the women in this study did not have clinical BV, 108 of them did report experiencing at least one risk symptom associated with BV within the 60 days prior to sample collection (108/285, 38.9%). Vaginal yeast infections were reported by 19 of the 285 respondents during the same time period. Most risk symptoms clustered together within the BN, with edges connecting vaginal discharge to both vaginal irritation and vaginal itch; a fishy/musty vaginal odor to staining of the underwear; any vaginal odor to vaginal itch; vaginal itch to vaginal infection; and vaginal infection to yeast infection specifically. Staining of the underwear was also directly connected to uro-abdominal pain in the preceding 60 days, which in turn was connected to self-treatment of a vaginal infection. While the Nugent score and vaginal pH were not connected to these risk symptoms, there were direct connections with specific microbial taxa. Presence of *Lactobacillus jensenii* increased the likelihood of a woman reporting uro-abdominal pain within the preceding 60 days, from a probability of 3.6% to 10.1% conditional on absence or presence of *L. jensenii*, respectively. Interestingly, PERMANOVA analysis showed an association between underwear staining in the previous 60 days and the vaginal microbiome (Table 1); the results of BN analysis suggest that uro-abdominal pain and *L. jensenii* may be the specific factors that link underwear staining and the microbiome. Use of pantyliners during menstruation was also associated with *L. jensenii*, with the conditional probability of *L. jensenii* being present in the vaginal microbiome increasing from 37.4% to 60.7% with use of panty liners during menstruation. While the directionality of this relationship is not immediately intuitive, previous work has shown that use of emollient pads changes the vaginal epithelium and that some women’s vaginal microflora does shift with pad versus tampon use (63,64), although the evidence on this is mixed (65). NMDS/ANOSIM testing indicated that tampon use was associated with the vaginal microbiome, lending further support to the hypothesis that menstrual habits could impact the microbiome (S5 and S6 Files). Alternatively, the support for edge directionality in the Bayesian network was based on majority rule and therefore directionality of arcs could potentially be reversed; in such a scenario, presence of *L. jensenii* could increase the likelihood of uro-abdominal pain, and thus underwear staining and pantyliner use. Furthermore, ethnicity was a potential confounder in the relationship between the microbiome and menstrual habits, as the network showed an influence of ethnicity on pad versus tampon use, which in turn directly affected pantyliner use (Fig. 4). The relationships between uro-abdominal pain, underwear staining, pantyliner use and *L. jensenii* deserve closer study given the importance of this microbe in vaginal health.

Other connections between bacteria and risk symptoms of itch, irritation, odor, staining, pain or yeast infection included use of a pad outside of menstruation increasing the conditional likelihood of a woman harboring *Proteobacteria* from 2.8% to 6.7%, as extra-menstrual pad use could be considered a proxy for vaginal discharge, staining or odor. Because *Proteobacteria* is such a diverse phylum that includes pathogens and non-pathogens, it is difficult to formulate hypotheses concerning these relationships. *Corynebacterium* seemed to play an important role in symptomology and sexual behaviors, as it was directly connected to vaginal odor and number of sexual partners in the 60 days preceding the survey. Women with this bacterium in their vaginal flora were more likely to have reported both no or multiple sexual partners, and were less likely to have reported a single sexual partner, while presence of *Corynebacterium* increased the risk of vaginal odor in the 60 days preceding the survey. The association of *Corynebacterium* with vaginal pathology is not widely described in the literature, although it has been recognized as an important vaginal community member for several decades (66). More recently, studies indicate that this bacterium is more prevalent in women with spinal cord injuries, women with treatment-refractory BV, and women with urgency urinary incontinence (67–69). Depending on the species, *Corynebacterium* can be fermentative, and many species produce amino acids, which can alter pH and the metabolite profile of the surrounding environment, thus providing a potential mechanism of *Corynebacterium’s* role in vaginal odor and pathology. These associations were not and would not have been easily observed using NMDS/ANOSIM and PERMANOVA.

Presence of *Bacteroides* had a protective effect against vaginal discharge; the probability of reporting discharge given the presence of *Bacteroides* was 2.0%, compared to 28.9% given absence of these bacteria within the vaginal microbiome. This runs counter to current notions about *Bacteroides*, an anerobic bacteria which is considered a commensal of the gastrointestinal tract and has been implicated in BV (70,71). However, not all vaginal discharge should be considered abnormal (72,73), and therefore it would not be a contradiction for *Bacteroides* to be associated with a decreased prevalence of “healthy” vaginal discharge (such as cervical mucous released during ovulation) and an increased prevalence of clinical BV. Distinctions such as these will become increasingly important as the scientific community attempts to attain a more nuanced understanding of connections between the microbiome and true clinical disease.

### Microbiome-Behavior Connections

Sexual behavior, sanitary practices and microbiome composition have all been associated with BV as separate factors. However, little is known about how a woman’s sexual and sanitary habits may influence the microbiome, or vice versa; and how this interplay may influence the likelihood of developing BV. The network resulting from this dataset showed several direct connections between the study participants’ habits and the microbiome. Whether or not women had ever had a pap smear was directly related to presence/absence of *Ureaplasma*. Women who reported having had a pap demonstrated a 34.1% likelihood of harboring *Ureaplasma*, compared to 18.0% among women who had never had a pap smear. Having had a Pap test was the only healthcare access question included in the Bayesian network that could be considered a proxy for socioeconomic status (48).

### Microbiome-Birth Control Interactions

Use of certain types of birth control could influence the microbiome by modulating hormone levels or by introducing chemical or physical barriers into the vaginal environment. Use of an intrauterine device (IUD) directly influenced the presence/absence of two different bacteria, namely *Lactobacillus gasseri* and *Sneathia*; in both cases, use of an IUD decreased the likelihood of finding the bacterium within the vaginal microbiome. Previous research on the effects of IUD use on vaginal microflora has been mixed, with both copper and hormone IUDs associated with higher levels of *Candida* fungi (74) but no associations with any bacterial alterations (74–76). Given the location of IUD use as a relatively highly-connected intercategory node in the Bayesian network, as well as its potential association with nonspecific vaginitis and recurrent BV (32,38,68), the associations with *L. gasseri* and *Sneathia* warrant follow-up study.

Women who reported ever using any form of birth control were more likely to have *Lactobacillus crispatus* within their microbiome compared to women who had never used birth control (conditional probability of 56.6% versus 48.1%, respectively). Women who reported male condom use in the 60 days preceding the survey were less likely to harbor *Lachnospiraceae* than women who did not (conditional probability of 1.3% versus 9.9%). Interestingly, PERMANOVA testing also showed a connection between male condom use in the previous 60 days and the overall composition of the vaginal microbiome (Table 1); the BN analysis allowed us to pinpoint this relationship to *Lachnospiraceae* specifically. As with other associations, there were some directed edges that did not immediately fit the paradigm for host-microbe interactions. For example, IUD use and abstinence were influenced by status of *Parvimonas* and *L. vaginalis* within the vagina, respectively. It is difficult to hypothesize a mechanism by which vaginal bacteria could influence a woman’s birth control decisions, even though recent studies have challenged the paradigm that host-microbiome interactions are largely uni-directional by suggesting that microbiome composition can directly influence host behavior (77–79). This highlights the importance of remembering that the directionality of edges in the BN are not definitive.

### Microbiome-Microbiome Interactions

Within the entire network, bacterial nodes fell largely within 2 modularity-based groupings, with some exceptions (Fig. 4). Upon isolation of bacterial nodes in a separate graph and subsequent modularity optimization, four communities of bacteria became evident (Fig. 7). These groupings did not seem to segregate by either Gram stain, cellular metabolism, or any other known characteristics. These groupings could be driven by as-yet undiscovered microbial relationships within the vaginal environment and/or by unmeasured, non-microbiome factors in this dataset.

**Figure 7.**
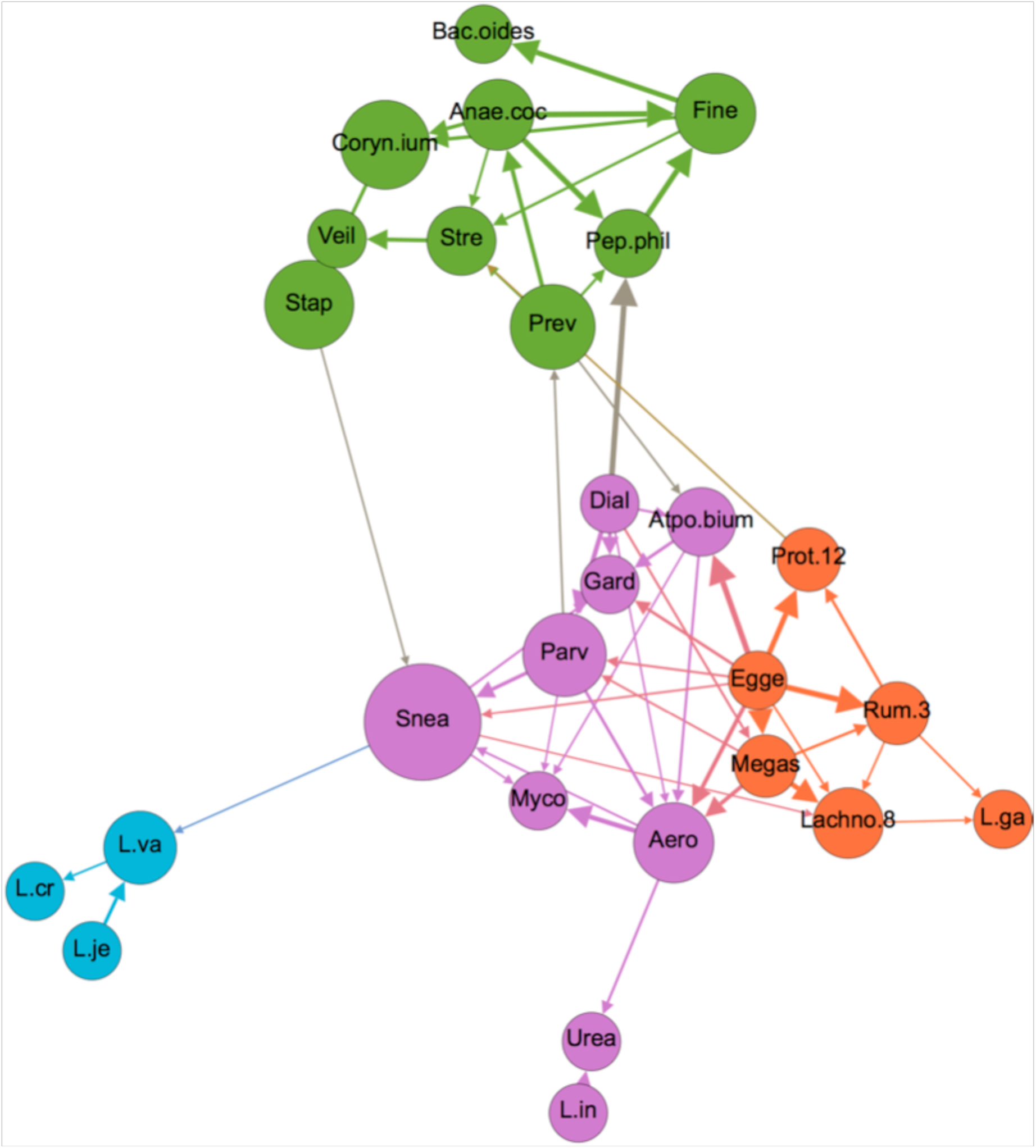
Network depicting only microbiome nodes, colored by modularity-based community type, as determined using a modularity optimizing algorithm (41). Node size is proportional to betweenness centrality, and edge thickness is proportional to bootstrap support. Node label abbreviations can be found in S4 File.

With the exception of *L. iners* and *L. gasseri*, the *Lactobacilli* formed a tight cluster to themselves. The fact that *L. iners* segregated separately from the other *Lactobacilli* is interesting given that this organism differs substantially in terms of genomic and metabolic characteristics, and its role in vaginal health has been debated despite its very high prevalence (47,80). One clue in this debate could be the tight association between *L. iners* and *Ureaplasma*, the latter of which has been implicated in several female reproductive conditions including chorioamnionitis leading to premature delivery and pelvic inflammatory disease (81,82). Presence of *L. iners* increased the likelihood of *Ureaplasma* from 19.4% to 36.3%, a significant increase that could have health implications and therefore deserves follow-up study.

## Conclusions

Using widely accessible software, we have demonstrated the utility of applying a Bayesian Network approach to a multi-dimensional microbiome dataset. Using this approach, we have demonstrated associations between women’s sexual and menstrual habits, demographics, vaginal microbiome composition and symptoms and diagnostics of BV. Many of these associations suggested intriguing relationships, indicating that the BN approach is able to highlight important associations within complex datasets, which can then be used for hypothesis generation. While follow-up studies are needed to investigate the significance of these novel associations, the validity of the associations was buttressed by the presence of many well-documented and self-evident connections within the overall BN. For instance, our BN confirmed the importance of vaginal pH and *Gardnerella* as influencers on the Nugent Score (Fig. 4). In addition, we found a very strong association between previous pregnancy, vaginal birth and tubal ligation. Indeed, all 17 participants who reported tubal ligation had previously been pregnant. The most recent data available suggest that nearly half of tubal ligations were performed in the postpartum period and that younger women were more likely to have tubal ligation performed post-partum (83). As with the BN produced here, this suggests a tight association between tubal ligation and previous pregnancy. Another confirmatory association showed that women who reported vaginal itch in the 60 days preceding the survey were much more likely to also have reported a yeast infection in the same time period. Vaginal itch has been closely associated with vulvovaginitis candidiasis (yeast infection) (84). While relationships such as these do not provide novel insight, they can be used as a “sanity check” in order to gain confidence in the structure of the BN to then move on to analysis of less well-understood network connections.

An advantage of the BN approach to datasets such as the one presented here is that BN algorithms are now easily accessible through well-documented and well-supported packages such as bnlearn (29), which has been integrated with the packages “parallel” (which is included in R-core) and “snow” (85) to support multi-threading for construction of large networks, and which now supports both categorical and continuous variables. Numerous GUI-based packages exist to support network visualization once the BN has been constructed (42,86), allowing for interactive graph exploration in an intuitive interface. Given the increasing use of multi-level, multivariate microbiome datasets, this accessibility should pave the way for more scientists to implement the BN analysis for such data. The need for such an approach has been demonstrated in the complexity of the relationships that we found within and between the vaginal microbiome, women’s sexual and menstrual habits, demographics and symptoms/diagnostics. While non-network approaches such as PERMANOVA and ordination/clustering can be used to uncover associations between individual metadata variables and the microbiome as a whole (as we demonstrate in this work), there are important limitations to these methods: first, these methods collapse the microbiome data into a single datapoint projected into two or a maximum of three dimensions, and therefore preclude assessment of intra-microbiome interactions, as well as interactions between metadata variables and specific microbiome members; second, these methods usually approach statistical testing on a variable-by-variable basis, and are thus ill-suited to understanding complex interactions between subsets of variables and the microbiome; and third, these approaches fail to show intra-metadata interactionswithin an analysis. BN analysis overcomes these limitations by testing for all possible associations between every node within the data – thus uncovering complex interactions within hierarchical, multidimensional datasets. Such datasets increasingly typify microbiome studies, and the need for rich and standardized metadata to support microbiome studies has been recently noted and implemented (87,88); such initiatives would be well-complemented by the data-flexible and intuitive nature of BN's. It is important to note, however, that the incorporation of more metadata variables (nodes) into BN analyses will necessitate larger sample sizes in order to support BN construction and inference. Similarly, the data distributions currently accommodated by available BN analysis packages for continuous variables are limited to the Normal (Gaussian) distribution. This constraint results in the need to discretize most microbiome data, which are commonly characterized by zero-inflated, skewed counts that do not conform to a Gaussian distribution. As was the case in this report, the need to dichotomize may cause loss of information, resulting in less-than-optimally refined inference. Fortunately, BN analysis methods form an active area of research, and extensions to accommodate distributions other than the Gaussian are currently being developed (89). As these efforts evolve, the BN approach is likely to become more flexible, and thus even better-suited to the unique characteristics of microbiome datasets.

## Acknowledgements

We would like to thank the editor and three anonymous reviewers for there useful comments.

## Supporting information

S1 File. Unfiltered, non-normalized, non-dichotomized microbiome counts and survey responses (upon request)

S2 File. Survey questions and abbreviations

S3 File. Filtered, discretized microbiome counts and survey responses for all 60 variables used in BN construction and analysis.

S4 File. *A priori* classifications, abbreviations, and BN analysis results (degree, closeness centrality and betweenness centrality) for all 60 variables used in BN construction and analysis.

S5 File. NMDS-ANOSIM testing results.

S6 File. NMDS plots of the six variables found to significantly influence the vaginal microbiome using the ANOSIM test.

**S7 Figure.**
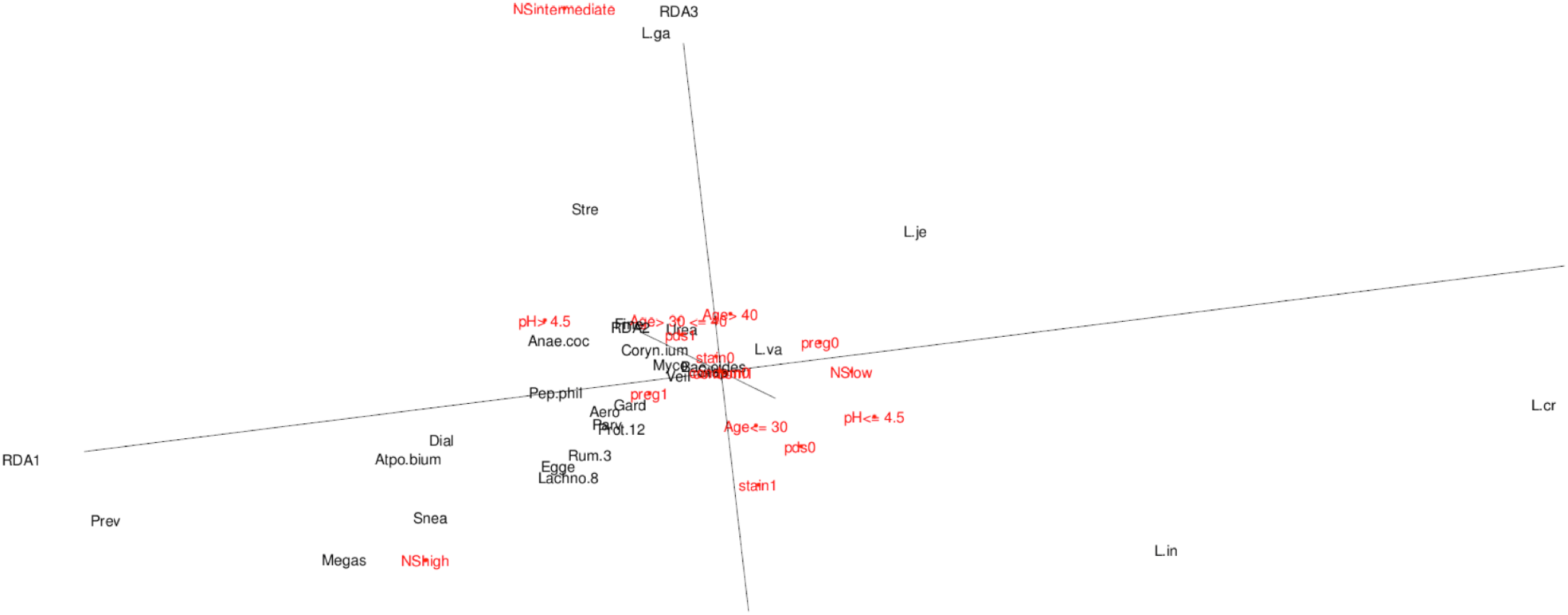
3D ordination plot based on PERMANOVA results.

S8 File. Figures describing the distribution of the abundances of taxa used in BN analysis.

